# GrapHiC: An integrative graph based approach for imputing missing Hi-C reads

**DOI:** 10.1101/2022.10.19.512942

**Authors:** Ghulam Murtaza, Justin Wagner, Justin M. Zook, Ritambhara Singh

## Abstract

Hi-C experiments allow researchers to study and understand the 3D genome organization and its regulatory function. Unfortunately, sequencing costs and technical constraints severely restrict access to high-quality Hi-C data for many cell types. Existing frameworks rely on a sparse Hi-C dataset or cheaper-to-acquire ChIP-seq data to predict Hi-C contact maps with high read coverage. However, these methods fail to generalize to sparse or cross-cell-type inputs because they do not account for the contributions of epigenomic features or the impact of the structural neighborhood in predicting Hi-C reads. We propose GrapHiC, which combines Hi-C and ChIP-seq in a graph representation, allowing more accurate embedding of structural and epigenomic features. Each node represents a binned genomic region, and we assign edge weights using the observed Hi-C reads. Additionally, we embed ChIP-seq and relative positional information as node attributes, allowing our representation to capture structural neighborhoods and the contributions of proteins and their modifications for predicting Hi-C reads. Our evaluations show that GrapHiC generalizes better than the current state-of-the-art on cross-cell-type settings and sparse Hi-C inputs. Moreover, we can utilize our framework to impute Hi-C reads even when no Hi-C contact map is available, thus making high-quality Hi-C data more accessible for many cell types.

**Availability:** https://github.com/rsinghlab/GrapHiC

**ACM Reference Format:** Ghulam Murtaza, Justin Wagner, Justin M. Zook, and Ritambhara Singh. 2018. GrapHiC: An integrative graph based approach for imputing missing Hi-C reads. In *Proceedings of 22nd International Workshop on Data Mining in Bioinformatics (BioKDD ‘23)*. ACM, New York, NY, USA, 16 pages. https://doi.org/XXXXXXX.XXXXXXX

## 1 Introduction

Research on genome organization has established its role in gene expression regulation [24], and how perturbations in this organization can lead to disease onset [13]. Hi-C, a high-throughput chromosome conformation capture experiment, allows researchers to understand and study genome organization. The Hi-C experiment produces an array of paired-end reads, representing the 3D structure of the genome (or chromatin) shown as an input to our pipeline in Fig. 1. Each paired-end read contains sequences of DNA interacting in the 3D space. A common way to aggregate the data produced by the Hi-C experiment is to store it in a contact map of size *N* × *N* . Each row and column correspond to fixed-width *N* windows (“bins”) in the range of 1 Kbp to 1 Mbp depending on the number of reads tiled along the genomic axis. The values in the contact map are counts of read pairs that fall into the corresponding bins. The study of these contact maps has revealed important structural features such as topologically associated domains (TADs) [5] and enhancer-promoter interactions [30] that are involved in gene regulation. Therefore, Hi-C experiments have proven to be crucial in helping us understand the interplay of spatial structure and gene regulation.

**Figure 1.**
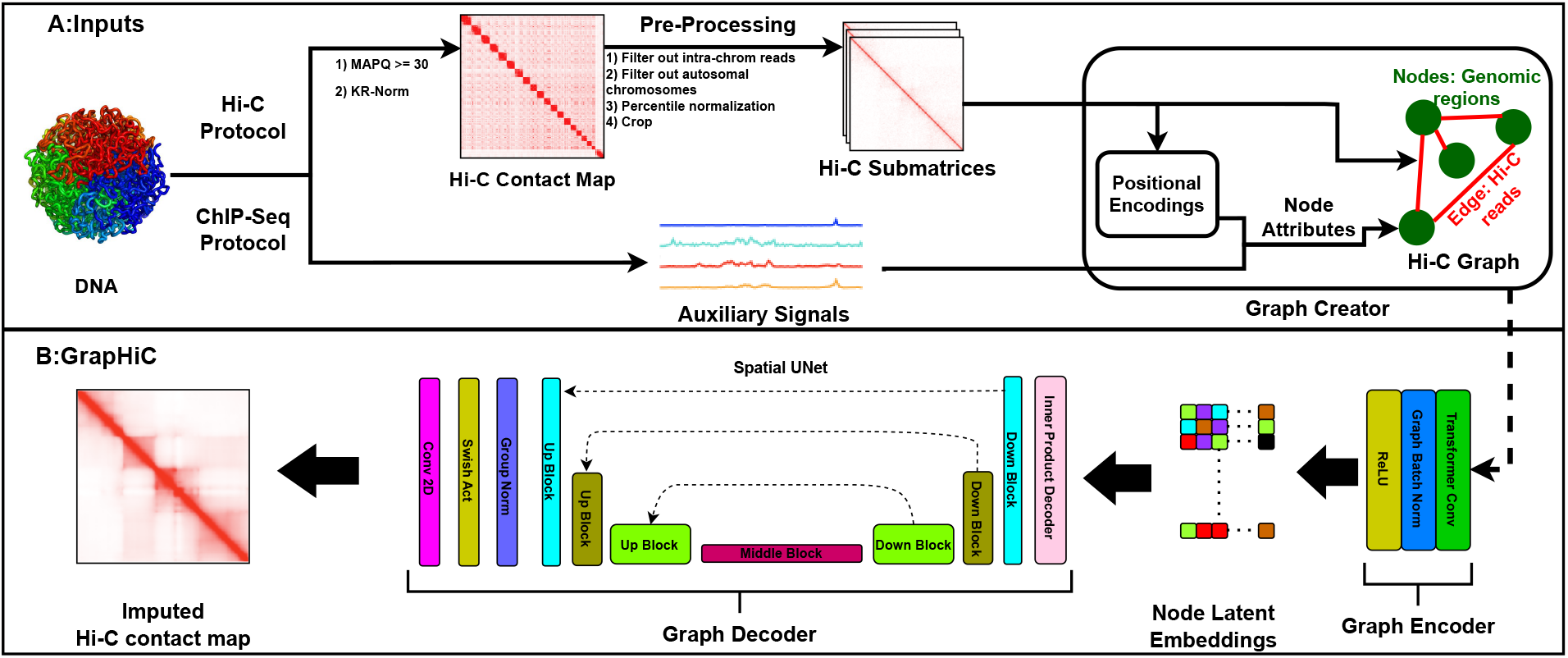
**A** Inputs and pre-processing pipeline: In this portion of the pipeline, we normalize Hi-C contact maps and create a Hi-C graph using Hi-C data as its edges and genomic loci as the nodes, with auxiliary genomic signals and positional encodings as node attributes. **B** GrapHiC: we use the sparse input Hi-C graph to generate a denser Hi-C graph. The Graph encoder takes in the Hi-C graph and generates latent node representations that our graph decoder utilizes to impute a dense Hi-C contact matrix.

Unfortunately, due to the quadratic scaling of reads in the Hi-C protocol, most experiments produce sparse read counts that require a larger bin size (typically more than ≥40 Kbp [37]) to account for the experimental noise. Consequently, any downstream analysis on such maps misses out on the finer structural features, such as enhancer-promoter interactions that typically occur in the 5 Kbp to 10 Kbp range [30]. Constructing Hi-C contact maps with sufficient resolution often requires billions of reads [30], which is often infeasible. This technical limitation of the Hi-C protocol restricts the thorough analysis of the 3D conformation of DNA.

To make Hi-C analysis more accessible, researchers have proposed several methods to impute the value of undetected reads in the Hi-C protocol. These methods can be classified into two categories; the first set of methods, HiC-to-HiC, uses a sparse Hi-C contact map and imputes the missing Hi-C reads by formulating it as an image resolution improvement task [11, 12, 18–20, 37]. The second set of methods, Seq-to-HiC uses cheaper-to-acquire data modalities such as DNA sequence [10], ChIP-seq signals [36] or a combination of both [33] to impute Hi-C contact maps. While Seq-to-HiC methods have fine-grained information about the proteins and their modifications that are known to mediate genome organization, they are inherently limited to not account for structural features such as A/B compartments[17] and TADs[5] that are also crucial in mediating interactions. HiC-to-HiC methods, on the other hand, implicitly rely on these structures to be available in the input contact map. However, they struggle to impute Hi-C reads when the contact maps are too sparse, and the structure is degraded [25].

We propose GrapHiC that combines both Hi-C and ChIP-seq data in a single graph-based representation to overcome both limitations. We formulate Hi-C data as a graph with nodes representing genomic loci, and the observed Hi-C reads as an edge (with weights) between them. We embed ChIP-seq signals and relative graph positional encodings as the node’s input features (or attributes) as summarized in Fig. 1 **A**. The positional encodings intuitively arrange nodes with the same structural features closer, such as TADs or A/B compartments, as node attributes. For our GrapHiC architecture, we implement a generative graph-based autoencoder that first encodes the input graph into a latent representation. This latent representation encodes the likelihood of two nodes interacting in the genome conditional on which TAD and A/B compartment they reside in, their structural neighborhood (edge weight), and the proteins/epigenome mediating the structure around them (via the 1D signals as node attributes). Then the graph decoder applies a UNet [26] on this latent representation to impute a denser Hi-C contact map.

In this work, we make the following key contributions:

1. We propose an end-to-end graph generative framework that outperforms the existing state-of-the-art methods by 21% on average (using a Hi-C similarity metric) when provided with five different sparse GM12878 inputs and 14% on average in cross-cell-type inputs.
2. We show the value of adding relative graph positional encodings in the Hi-C graph formulation through our ablation analysis. Positional encodings improved our performance by 24% over a Graph formulation without positional encodings.
3. We provide a proof-of-concept result demonstrating that we can use a Hi-C map with expected reads (whose probability decays exponentially with distance from the diagonal) with ChIP-seq data to impute high-quality Hi-C contact maps reliably when a low-quality Hi-C input is unavailable.

## 2 Related Work

Existing HiC-to-HiC methods use low-coverage^1^ Hi-C contact maps to impute high-coverage (or high-read-count) outputs. These methods formulate this imputation as an image resolution improvement task [6] by treating the Hi-C contact maps as images. Convolutional Neural Networks (CNNs) extract a feature set from the input low-read-count contact map and then use those features to predict a high-read-count contact map. HiCPlus [37] utilized a 3-layer CNN and optimized this model using a Mean Squared Error (MSE) loss. Later methods extended over HiCPlus by stacking more layers [19, 20], employing Generative Adversarial Networks (GAN) style training [12, 18] or by using nuanced biologically-grounded loss functions [11] to impute more realistic contact maps.

On the other hand, existing Seq-to-HiC methods use cheaper-to-conduct 1D genomic signals as inputs, such as DNA [10], ChIP-seq [36] or a combination of both [33], to predict Hi-C reads using decision trees [36] or deep learning based approaches [10, 33]. Even though Seq-to-HiC methods have fine-grained positions of proteins (or their modifications) that are known to mediate the genome conformation, they miss out on the structural neighborhood. While HiC-to-HiC methods capture this structural neighborhood, they struggle to generalize when the input Hi-C contact map becomes too sparse and the structure degrades significantly [25]. To overcome both limitations, we propose combining low-read-count Hi-C contact maps and the cheaper-to-conduct ChIP-seq signals in a single graph-based representation.

Other related efforts have also suggested formulating Hi-C data as a graph. For example, Hi-C graph formulation has been previously used for gene expression prediction [2], chromatin state prediction [15], Micro-C prediction [8], and many other tasks including computing similarity between two Hi-C contact maps [35]. Our method resembles Caesar [8], but we make a few critical methodological innovations and changes. First, Caesar applies an ensemble model that predicts the Micro-C contact map and Chromatin Loops independently of each other, while we propose to utilize a single model. We believe our formulation allows our model to learn robust internal representations because both tasks rely on the same underlying features. Secondly, we use a UNet architecture to decode the Hi-C contact map because a UNet architecturally mimics the hierarchical organization of the chromatin and can potentially learn better internal representations. Lastly, we use eigenvectors of the Laplacian of our Hi-C graph as graph-positional encodings that, intuitively, positions the nodes belonging to the same graph communities, clusters, or structures closer together in the graph space. In the biological context, particularly the sign of the eigenvectors divides the genome into two A/B compartments, where genomic loci in A compartment are more likely to interact with each other and vice-versa [17], and these structures inform the global organization of the genome [30]. From a graph learning perspective, positional encodings are helpful in all tasks that formulate Hi-C as a graph because they expand the theoretical expressiveness of graph-convolutional operators beyond the 1-Weisfeiler-Lehman (1-WL^2^) isomorphism test, allowing GrapHiC to tell apart similar structures, such as TADs in the latent space based on their position in the chromatin [28].

## 3 Methods

Our method aims to impute a high-resolution Hi-C contact map *H*_*imputed*_ given a sparse Hi-C contact map *H*_*sparse*_ . We develop GrapHiC, shown in Fig. 1, which has three main components. First, we develop a **Graph creator** module, depicted in Fig. 1 **A**, that models Hi-C contact maps as graphs, with graph positional encodings and ChIP-seq signals as node attributes, and observed Hi-C reads as edges connecting those nodes. Second, we implement a Graph Autoencoder, shown in Fig. 1 **B**, that produces a dense Hi-C graph when provided with a sparse Hi-C input. The **Graph encoder** utilizes Graph Transformer convolutions that explicitly attend to both the node features and edge weights to learn latent node embeddings. Last, our **Graph decoder** uses these latent node embeddings to impute a dense Hi-C contact map *H*_*imputed*_ using a UNet architecture.

### 3.1 Graph Creator

Given a sparse Hi-C contact map *H*_*sparse*_, *Graph Creator* module defines a Hi-C graph *G* = (*X, E*), with a node attribute set *X* and an edge set *E* connecting those. Here, a node *i* ∈*X* corresponds to genomic loci and an edge *e*_*i,j*_ ∈*E* connecting them, provided through observed reads in our sparse contact map *H*_*sparse*_ . Most existing Hi-C graph formulations define *X* to be a constant set [9, 16] and more recently capture their genomic profiles [2, 8]. We believe this node attribute set *X* should also include the relative node position because, for example, loci that are further apart are less likely to interact [30]. In contrast, loci belonging to the same A/B chromatin organizational compartments or TADs are more likely to interact with each other [5, 17].

We compute the relative position of the nodes using the graph positional encoding scheme [7], that decomposes the Laplacian of the input sparse Hi-C contact map *H*_*sparse*_ into the spectral domain that provides us a set *N* eigenvectors *V*, where top *k* components of the eigenvector 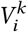 corresponds to the relative position of node *i* in the input Hi-C contact map domain. Intuitively, these positional encodings assign the node a unique position in the graph space with similar values to nodes belonging to the same sub-structures, such as TADs and A/B compartments. Lastly, to integrate the genomic information of a locus, we take the average reads for that region from five ChIP-Seq signals (DNAse-Seq, CTCF, H3K4ME3, H3K27ME3, H3K27AC) to define a feature set *C*. Finally, we concatenate *V* with *C* to get our node attribute set *X* .

### 3.2 Graph Encoder

The *Graph Encoder* constructs latent representations of each node that is a weighted aggregation of its features and the node features of its neighboring nodes. We use graph transformer convolutions [32] in the graph encoder. Graph transformer convolutions learn the new latent attribute set 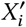 of the node *i*, 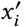, by aggregating over all *N* nodes with features *x*_*i*_ ∈ *X* as follows:

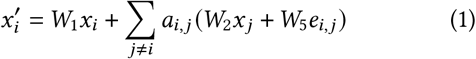

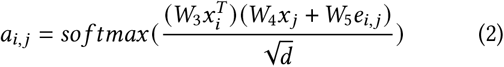

Here, *e*_*i,j*_ ∈*E* are edge weights connecting node *i* and *j, a*_*i,j*_ are attention coefficient attributes, *d* is a scaling parameter, and *W*_1_ to *W*_5_ are learnable parameters. *W*_5_ is shared in calculation of both the attention coefficients *a*_*i,j*_ and the new latent attributes set 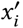. The graph transformer convolution operation updates the representation of the current node by combining the current node’s features *x*_*i*_ with node features of neighboring nodes *j* ≠ *i* and edge weight connecting them *e*_*i,j*_ (Eq. (1)). This combination is scaled by their attention coefficient *a*_*i,j*_ (Eq. (2)). Attention allows the model to focus on the most relevant node interactions by weighing them differently, with weights ranging between 0 - 1 due to the *so f tmax* function. In the biological context, this latent representation encodes the likelihood of two nodes given their structural neighborhood (encoded through relative positioning) and the genomic landscape mediating the structure around them.

### 3.3 Graph Decoder

*Graph Decoder* module starts off by taking the inner product of the latent node embeddings *X*^′^ with the transpose of themselves to get a contact probability map *P* that is of shape *N* × *N* as follows:

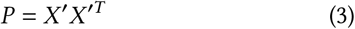

*Graph Decoder* then applies a UNet [26] to transform the contact probability map into observed Hi-C reads contact map by first extracting a feature set 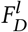 using 3 *Down Blocks*, that work by applying Downsampling convolutions. At the *Middle Block* we transform the feature set 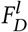 to 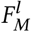 by applying self attention:

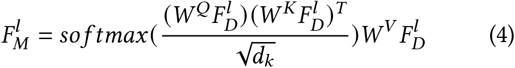

Here *W* ^*Q*^, *W* ^*K*^, *W* ^*V*^ are learnable parameters and *d*_*k*_ is a scaling parameter. This self-attention operation in a biological context learns the relationships between how the properties of sub-structures such as sub-TADs relate to other sub-TADs in the neighborhood and the overall structure of the genome. Next, UNet uses this feature set 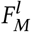 to impute a dense Hi-C contact map by applying 3 *Up Blocks* that use Deconvolutions (that are inverse of convolutions in *Down Blocks*) to impute a Hi-C contact map. *Up Blocks* in comparison to *Down Blocks* have an additional cross-attention operation that learns the relationships from the feature set from the previous *Up Block* (or *Middle Block*) and the features from the same level *Down Block* highlighted as dotted gray connections in the Fig. 1. This cross-attention operation in a biological context allows higher-order chromatin organization features extracted in the upper level *Down Blocks*, such as A/B compartments, that might get lost in the bottleneck *Middle Block* to inform Hi-C read imputations. Lastly, *Graph Decoder* applies a group norm, swish activation [12], and last convolution operation to project the output of the last *Up Block* to our final imputed Hi-C contact map *H*_*imputed*_ .

We jointly optimize both the Graph Encoder and the Graph Decoder, GrapHiC, with MSE loss as follows:

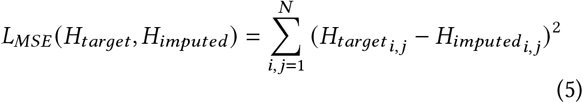

We selected MSE loss because it has been shown [14] that having complex, nuanced loss functions increases the likelihood of the model overfitting to the training objective. This problem is more relevant in the graph domain because graph convolutions tend to overfit the underlying geometry [23].

### 3.4 Implementation Details

We use an ADAM optimizer with a 0.0001 learning rate to optimize both the encoder and the decoder. We implement the entire pipeline in Python (version 3.9.0), Pytorch (2.0.0), and Pytorch Geometric. We show the details of the model architecture in Supplementary Table S1. GrapHiC takes in a 256 × 256 sized Hi-C sub-matrix corresponding to 2.56 Mpb regions (because of the 10 Kbp resolution). GrapHiC also requires a 256 × 5 ChIP-seq profile of the same genomic region to produce a Hi-C graph *G* with 256 nodes, and each node has an attribute vector of size 13 (5 ChIP-seq signals and top 8 components of the eigenvectors). The size of 256 serves two purposes. First, it allows us to include all the biologically informative interactions in the 2 Mbp range around the diagonal. Second, it ensures that we can downsample by a factor of 2 three times (number of Down Blocks) in our UNet architecture. To predict intra-chromosomal contact maps (similar to our related works [8, 11, 12, 19, 33, 37]), we predict along the diagonal by sampling 2.56 Mbp sub-matrices with a 0.3 Mbp stride length. We average all the overlapping predictions to account for the border effect and produce our final intra-chromosomal contact map. We trained GrapHiC with 10 random seeds and found a standard deviation of 0.004 on validation chromosomes on SSIM (mentioned in the next section). We trained GrapHiC on another random seed and used that for the rest of the evaluations.

## 4 Experimental Setup

### 4.1 Datasets and pre-processing

Following existing Hi-C read imputation methods [11, 12, 18–20, 37], we used GM12878, IMR90, and K562 cell line datasets from Rao *et al*. [30] as our target high-read-count (HRC) matrices. Given the technical constraints in acquiring and reprocessing (both Hi-C and ChIP-seq data), we leave cross-species analysis as future work to ensure the data distributions match. These are the ground-truth high-quality datasets in our experiments that we would like GrapHiC to impute using the low-quality Hi-C contact maps as inputs (as done in previous works). As summarized in Table. 1 we collected eight low-read-count (LRC) Hi-C contact maps from the ENCODE [21] and the NCBI [1] public repositories with read coverage in the range of 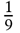 to 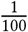 of the reads in comparison to the appropriate HRC Hi-C contact maps. We also include a dataset from the 4DN portal for additional evaluations on GRCh38 (for additional results) shown in the Supplementary Table. **??**. We removed spurious and incorrectly mapped reads by applying a MAPQ filter of ≥30. We binned the remaining reads at 10 Kbp resolution (size of genomic loci) to create our two-dimensional contact maps. We performed KR-normalization of these contact maps to balance reads across all the bins. Moreover, we filtered out all inter-chromosomal contacts and confined our analysis to autosomal chromosomes. We normalized each contact map between 0 and 99.9^*th*^ percentile values following DeepHiC’s normalization procedure.

**Table 1.** Summary of the datasets and their sources. HRC refers to high-read-count, and LRC refers to low-read-count Hi-C contact matrices. Sparsity represents the fraction of reads compared to the relevant HRC Hi-C contact map.

|  | Reads | Sparsity | Source |
| --- | --- | --- | --- |
| **GM12878-HRC** | 1,844,107,778 | 1 | GSE63525 |
| **IMR90-HRC** | 735,043,093 | 1 | GSE63525 |
| **K562-HRC** | 641,402,880 | 1 | GSE63525 |
| **GM12878-LRC-1** | 202,380,884 | 9 | GSE63525 |
| **GM12878-LRC-2** | 70,138,184 | 25 | ENCSR968KAY |
| **GM12878-LRC-3** | 42,453,795 | 44 | ENCSR382RFU |
| **GM12878-LRC-4** | 37,079,587 | 50 | ENCSR382RFU |
| **GM12878-LRC-5** | 18,696,952 | 100 | GSE63525 |
| **IMR90-LRC-1** | 75,193,876 | 10 | GSE63525 |
| **K562-LRC-1** | 44,882,605 | 14 | GSE63525 |

For our auxiliary genomic signals, we used 5 out of the 14 1D inputs used by HiCReg [36]. HiCReg curated a collection of ChIP-Seq experiments targeting ten histone marks (notably H3K27AC and H3K27ME3), three transcription factors (RAD-21, CTCF, and RNA-Pol2), and one chromatin accessibility marker (DNAse-Seq), essentially forming a comprehensive set of features related to the 3D organization of the DNA. Like our Hi-C data, we binned all the 1D data in 10 Kbp genomic bins, normalized them in the 0 to 99.9^*th*^ percentile range, and cropped them into sub-ranges of size 2 Mbp that aligned with Hi-C submatrices. As done in previous works, we divided chromosomes [chr1-chr8, chr12-chr18] as training, [chr8, chr10, chr19, chr22] as validation, and [chr9, chr11, chr20, chr21] as testing sets. All of our results are on cross-chromosome evaluations (on the testing set), at no point during the training procedure we use data from these chromosomes.

### 4.2 Baselines

We compare our method against two state-of-the-art Hi-C read imputation frameworks. We train baselines with all the pre-processed data (including normalizations) with the same pipelines we use for GrapHiC to ensure a fair comparison of our outputs.

#### HiCNN

A recent evaluation [25] showed that HiCNN [19] provides the best imputation generalizability across a broad range of sparse real-world Hi-C datasets in comparison to rest of the HiC-to-HiC imputation methods (including DeepHiC). HiCNN relies on a 54-layer CNN that takes in 40 × 40 sparse Hi-C contact map sub-matrices and predicts high-resolution Hi-C sub-matrices across 2 Mbp distance from the diagonal. HiCNN then assembles those sub-matrices into intra-chromosome contact maps.

#### HiCReg

We include HiCReg, a Seq-to-HiC baseline, that uses 14 ChIP-seq signals as input to impute Hi-C reads using a random-forest model. Similar to GrapHiC and HiCNN, we use HiCReg to predict intra-chromosomal contact maps in the 2 Mbp range along the diagonal to ensure a comparable output.

Note, we exclude Caesar [8] and Origami [33] from our baselines because we were unable to obtain the same quality Hi-C (or Micro-C) contact maps. Origami produced a Hi-C contact map with large regions with no observed reads, as shown in Supplementary Fig. S1 when provided with hg19 reference genome^3^ aligned DNA sequences, CTCF, and ATAC-Seq data. When provided with a real-world sparse Hi-C contact map, the matrices produced through Caesar (a Micro-C contact imputation framework) show severe degradation in genome structure compared to when we input a dense Hi-C contact map as depicted in Supplementary Fig. S2.

### 4.3 Evaluation Metrics

We compare the imputed Hi-C contact maps *H*_*imputed*_ against the target Hi-C contact maps *H*_*target*_ using the following three evaluation metrics:

**Structural Similarity Index Metric (SSIM)** compares the visual similarity of two images by comparing the luminance and contrast of small patches across the entire image to compute a score between 0 and 1, with 1 given to identical images. We use SSIM to compare the visual similarity of Hi-C contact maps similar to our related efforts[12].

**GenomeDISCO** [35] utilizes random graph walks of increasing lengths to compare the similarity of the underlying graph structural features. These graph structural features are known to be associated with higher dimensional chromatin features. GenomeDISCO produces a similarity score between -1 and 1, where the higher score represents higher similarity. We show GenomeDISCO scores because it computes similarity based on the structure rather than observed Hi-C reads distribution. Moreover, the random walk algorithm not only smooths out Hi-C protocol noise but also makes GenomeDISCO more robust to Hi-C read depth [35].

#### Chromatin Loops F1 score

To estimate the biological utility of our imputed chromatin maps, we compare the positions of the Chromatin Loops in the imputed Hi-C contact map against their position in the high-resolution Hi-C contact map. First, we call loops for both high-read-count matrices and imputed matrices using Chromosight [22], and then we compute F1 scores by counting: 1) True positives (TP) features that overlap in both matrices. 2) False Positives (FP), features that are called on the imputed matrices but are not present in the high-read-count matrices. 3) False Negatives (FN), features in high-read-count matrices that were absent in imputed matrices. Then we compute the F1 score using the following:

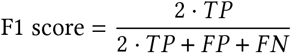

We include the other metrics in the Supplementary Section to benchmark our method thoroughly like HiCRep [35], Pearson Correlation Coefficient (PCC), and F1 scores for TAD boundaries (borders) and DNA-Hairpins. Hi-C-specific similarity metrics and feature recovery analysis metrics perform their own Hi-C normalizations and read downsampling procedures to ensure a fair comparison of Hi-C contact maps.

## 5 Results

### 5.1 Ablation analysis demonstrates the importance of positional encoding in graph formulation

We conducted an ablation analysis on our proposed graph formulation by comparing performance on the test chromosomes of the GM12878-LRC-3 dataset to investigate our design choices. We compare our imputed Hi-C contact maps against the GM12878-HRC-1 test chromosomes to evaluate performance. Here we compare three relevant versions of GrapHiC:

#### GrapHiC-Basic

The simplest graph formulation without any positional information or auxiliary ChIP-Seq experiments as node features. GrapHiC-Basic only uses the sparse Hi-C contact map embedded as edge weights to impute a Hi-C contact map.

#### GrapHiC-Pos

GrapHiC-Pos, similar to GrapHiC-Basic, also only takes in the sparse Hi-C contact map, but it generates graph positional encodings from the sparse contact map and embeds them as node attributes.

#### GrapHiC

Our default version takes in sparse Hi-C contact map and five ChIP-seq experiments CTCF, DNASE-Seq, H3K4ME3, H3K27AC, H3K27ME3. It generates graph positional encodings through sparse Hi-C contact maps and embeds them with ChIP-seq as node attributes.

Qualitatively, in Fig. 2(A), we compare the imputed Hi-C contact maps for the region chr11:20.1 Mbp-22.1 Mbp; we picked this region because it shows a high density of chromatin features, including TADs and Chromatin Loops ^4^. GrapHiC-Basic generates a contact map that resembles the expected contact map without any higher-order chromatin features. Although adding graph positional encodings in GrapHiC-Pos improves the structure we recover, adding auxiliary ChIP-Seq signals allows GrapHiC to recover the finer architectural features, particularly Chromatin Loops, as highlighted with a blue dotted rectangle. We quantify this improvement by showing the performance by comparing performance on three metrics (on the x-axis) SSIM, GenomeDISCO, and Chromatin Loops F1 score (on the y-axis) on test chromosomes of GM12878-LRC-3 dataset (training dataset) Fig. 2(B). Our results show that adding graph positional encodings improves SSIM and GenomeDISCO scores by 9% and 25% over the GrapHiC-Basic, respectively, highlighting the utility of relative positional encodings in recovering higher-order chromatin structure. Moreover, adding graph positional encodings improves Chromatin Loops recovery F1 scores by a substantial 88% showing their utility in recovering finer structural features. Adding ChIP-seq signals in the node attribute vectors marginally improves the SSIM and GenomeDISCO scores by 0.4% and 2.0%, respectively. However, adding five ChIP-seq signals, including CTCF, a protein known to mediate chromatin organization via forming chromatin loops by binding with cohesin [27], improves Chromatin Loops F1 score scores by an additional 20%. We perform further ablation studies, including how GrapHiC scales to the number of ChIP-seq signals, and validate our results across other datasets and metrics. GrapHiC’s scores are robust to the number of ChIP-seq signals across various metrics and datasets, as summarized in Supplementary Tables S4, S5, S6, S7, S8. This ablation analysis highlights the importance of including relative positional information as node attributes in formulating Hi-C as a graph and its utility in recovering biologically informative Hi-C contact maps. For the rest of our evaluations, we use the **GrapHiC** variant, which includes five ChIP-seq signals and relative positional information, to impute Hi-C contact maps unless stated otherwise.

**Figure 2.**
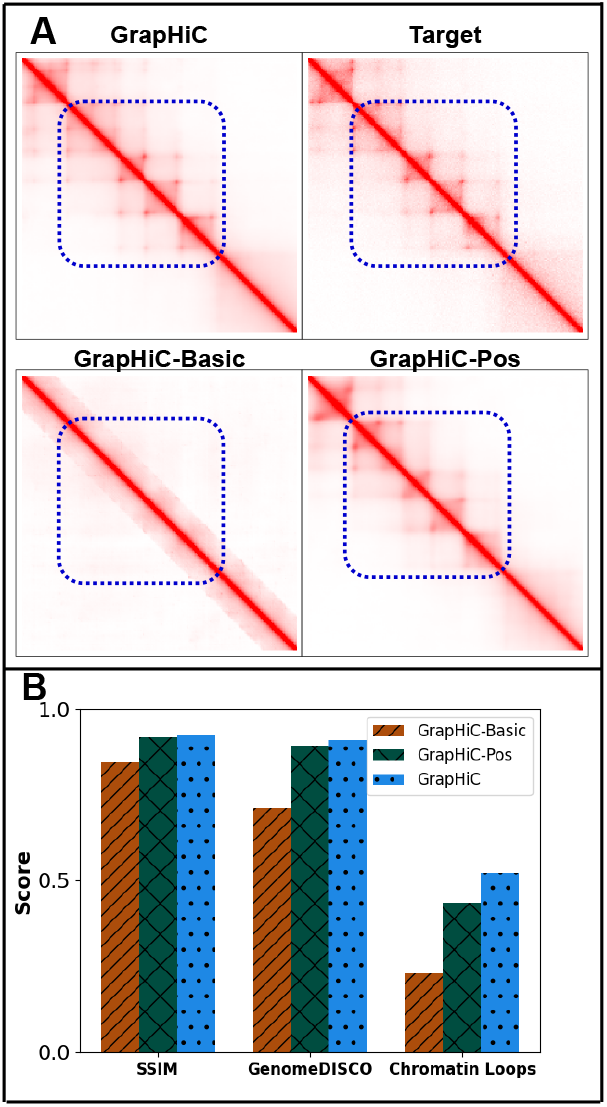
Positional Encodings improve the quality of imputed Hi-C reads. A visual comparison of the Hi-C contact map for the region chr11:20.1.5-22.1 Mbp generated by various versions of GrapHiC shows that as we add, relative positional encodings GrapHiC can impute a more realistic Hi-C contact map, and as we add ChIP-seq signals GrapHiC can recover finer architectural features better highlighted with a dotted blue rectangle. **B** Our quantitative analysis on SSIM and GenomeDISCO show similar scores for GrapHiC-Pos and GrapHiC, but Chromatin loop recall scores confirm our visual hypothesis suggesting that adding more ChIP-seq signals help GrapHiC to recover finer architectural features

### 5.2 GrapHiC outperforms existing methods on Hi-C datasets with varying levels of sparsity

We compare the performance of GrapHiC with HiCNN and HiCReg, which are HiC-to-HiC and Seq-to-HiC read imputation methods, respectively. We train both HiCNN and GrapHiC with GM12878-LRC-3 Hi-C contact map as input, and we test them both on the GM12878 cell line with five sparsity levels ranging from a 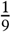 to a 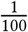 of the total reads in the target HiC map (GM12878-HRC-1). Note we show the same scores for HiCReg for all five GM12878 sparse inputs because HiCReg relies only on ChIP-seq data to impute missing reads and is agnostic to Hi-C reads sparsity.

As shown in Fig. 3 **A**, we visualize an imputed region chr11:20.1Mbp-22.1Mbp because it captures a cluster of TADs, sub-TADs, and Chromatin Loops. In the section highlighted with the blue dotted rectangle, we observe that GrapHiC can more accurately capture Chromatin Loops [29] compared to both HiCReg and HiCNN. To quantitatively evaluate these methods, we compare SSIM, GenomeDISCO, and chromatin loop F1 scores in Fig. 3 **B**. GrapHiC outperforms HiCNN by 11%, 21%, 12% and HiCReg (when comparing GrapHiC’s performance on GM12878 dataset 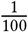 of reads) by 22%, 26%, 89% on SSIM, GenomeDISCO and Chromatin Loop F1 scores respectively. GrapHiC shows the highest performance improvement on the most sparse input Hi-C contact map (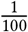 of reads) against HiCNN by 32%, 69% and 53% on SSIM, GenomeDISCO, and Chromatin Loops F1 scores highlighting GrapHiC’s capabilities to combine multiple modalities^5^ of data and impute biologically informative reads, especially when provided with a highly sparse Hi-C input. Our results on the other metrics in Supplementary Table S9 show similar trends that GrapHiC outperforms the baseline methods for the most sparse Hi-C datasets. However, HiCNN performs similarly or slightly better than GrapHiC for less sparse datasets (GM12878-LRC-2) because they have more genome structure in the input. This structure is also similar to the training samples, which supports HiCNN in good Hi-C read imputation and feature recovery. Based on these results, we conclude that GrapHiC can learn robust internal representations even when provided with 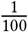 sparse (GM12878-LRC-5) input and can generalize to a wide range of sparse inputs.

**Figure 3.**
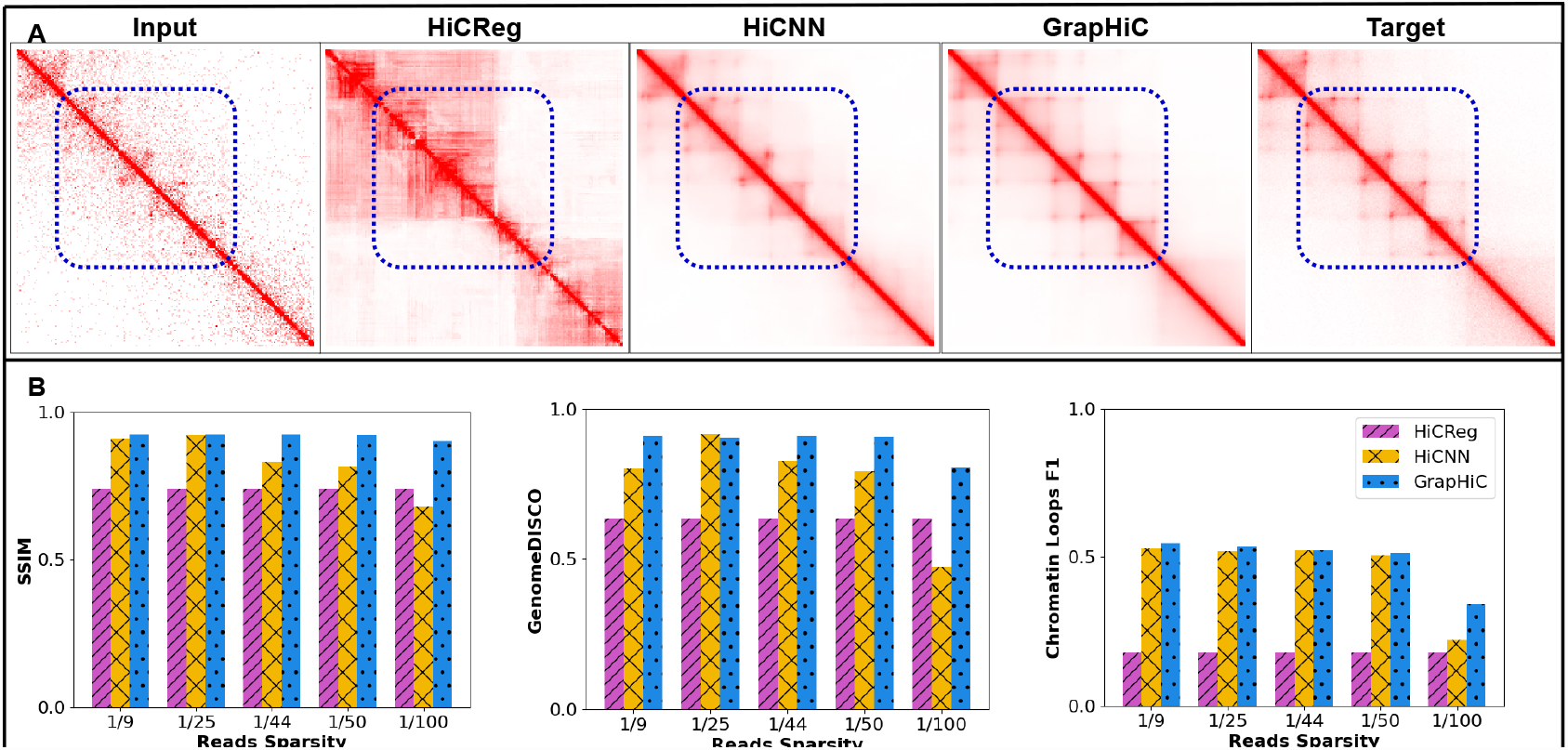
GrapHiC generalizes better to sparse GM12878 datasets. **A** Visual comparison of Hi-C contact map for the region chr11:20.1Mbp-22.1Mbp generated by HiCReg, HiCNN, and GrapHiC shows that GrapHiC can better recover finer chromatin architectural features highlighted with a dotted blue rectangle. **B** Our quantitative evaluation using SSIM, GenomeDISCO, and Chromatin Loops F1 scores (on the y-axis) suggests that GrapHiC outperforms the other methods in at least four out of five datasets across all metrics (on the x-axis) while showing most improvements in the sparsest input 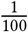 case.

### 5.3 GrapHiC trained on GM12878 generalizes well to IMR90 and K562

Next, we demonstrate that our GrapHiC model, trained on the GM12878 cell line, can impute the Hi-C maps of different cell lines. We input the LRC Hi-C contact maps and ChIP-seq signals for K562 and IMR90 and impute Hi-C contact maps using GrapHiC, HiCNN, and HiCReg.

First, we show the imputed Hi-C contact maps from region chr20:49.2-51.2 Mbp generated for IMR90 LRC Hi-C in Fig. 4 **A** and K562-LRC Hi-C in Fig. 4 **B** that have 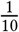 and 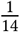 of the read coverage in comparison to their corresponding HRC Hi-C contact maps, respectively. GrapHiC imputes the most similar Hi-C contact map across both cell samples compared to the target HRC Hi-C contact map. Moreover, GrapHiC can recover cell-specific sub-TADs in the K562 sample and accurately predict its absence in the IMR90 sample, as highlighted with a blue rectangle in Fig. 4. Our quantitative analysis, shown in Fig. 4 **C**, on SSIM, GenomeDISCO, and Chromatin Loops F1 score shows that GrapHiC can generalize better to other cell lines than HiCNN and HiCReg. Specifically, GrapHiC improves SSIM scores by 19% and 27%, GenomeDISCO scores by 19% and 9%, and Chromatin Loops F1 scores by 61% and 26% on IMR90 and K562 cell lines, respectively, in comparison to HiCNN. Our results on other metrics (Supplementary Table S10) report that GrapHiC out-performs HiCNN on most metrics except HiCRep, QuASAR-Rep, and TAD recovery F1 scores, which are metrics that tend to assign more value to features that exist closer to the diagonal [5, 35]. Given there is a higher density of reads around the diagonal because of the distance effect in Hi-C protocol [3], HiCNN can impute Hi-C better reads in that region in comparison to GrapHiC that first maps the inputs into a latent representation and then impute Hi-C reads. Our qualitative and quantitative evaluations conclude that GrapHiC, trained on GM12878, can generalize to other cell lines substantially better than the baseline methods. Note that we observe a performance decrease compared to its performance on the GM12878 cell line datasets partly because GrapHiC generates reads that mimic GM12878’s higher sequencing depth and partly because of the significant distributional shift in Hi-C samples (particularly IMR90) between cell types [25]. Cell type tends to have a higher impact on graph-based formulations given their theoretical formulations forcing them to overfit to the underlying graph geometry [23].

**Figure 4.**
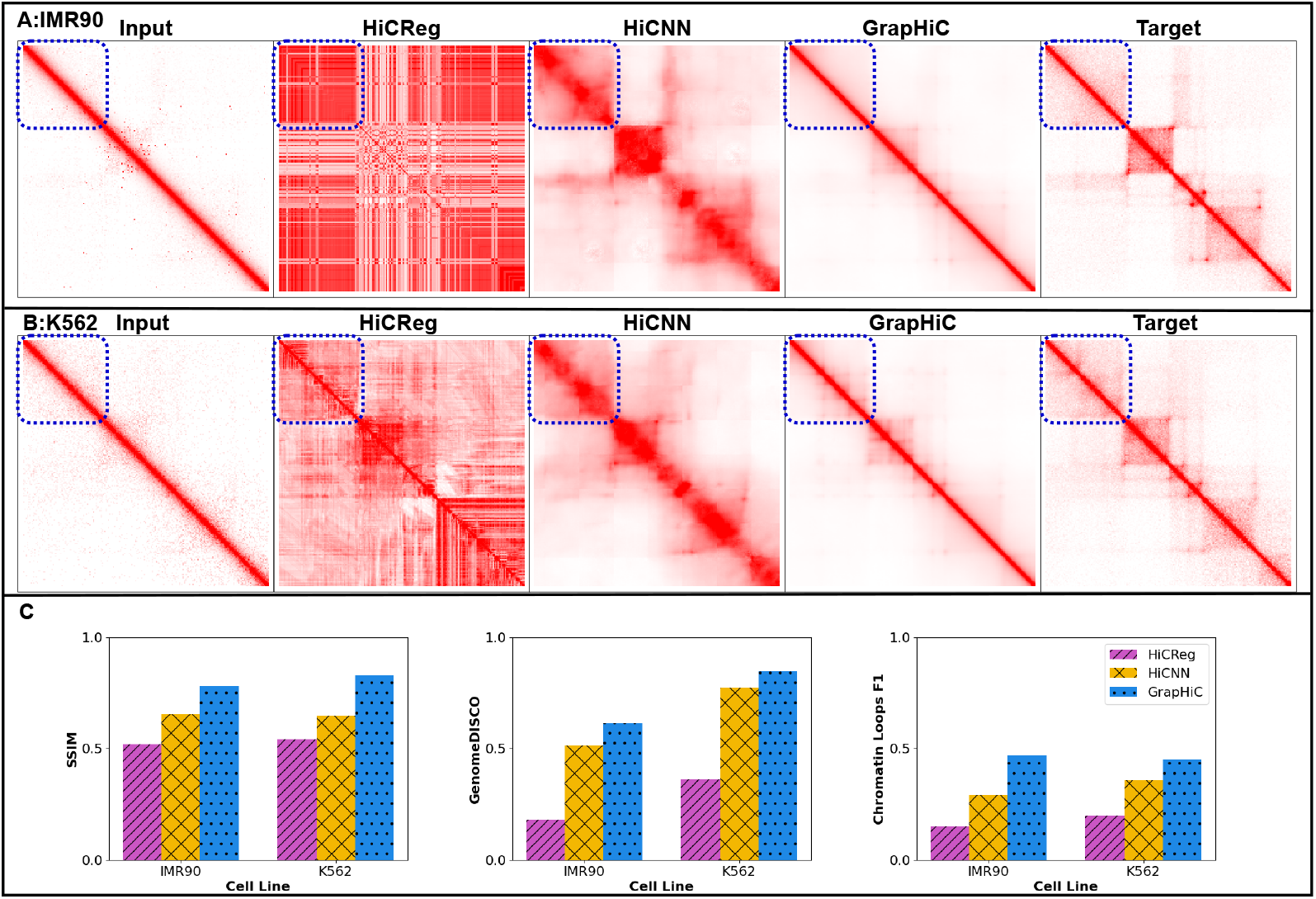
GrapHiC generalizes better than existing methods to IMR90 and K562 cell-lines. This figure shows the visual comparison of imputed Hi-C samples from GrapHiC, HiCNN, and HiCReg from IMR90-LRC-1 **A** and K562-LRC-1 **B** inputs for the region chr20:49.2Mbp-51.2Mbp to show cell-specific features. We show that GrapHiC cannot only impute a highly similar contact map but also recovers cell line-specific features highlighted with the dotted blue rectangle. **C** We compare the SSIM, GenomeDISCO, and Chromatin Loop F1 scores of GrapHiC, HiCNN, and HiCReg in a cross-cell type imputation scenario. We find that GrapHiC is able to generalize better than both HiCNN and HiCReg.

### 5.4 GrapHiC can produce high-fidelity Hi-C contact maps with missing Hi-C input

Given the unique multimodal nature of GrapHiC, we can impute Hi-C contact maps even in the absence of input LRC Hi-C data. Because our model learns representations based on multiple datasets it can potentially show resilience to the lack of one data type, for example, as we show in our ablations that we can impute high fidelity Hi-C contact maps without ChIP-Seq (GrapHiC-Pos). Note, this a task that HiC-to-HiC methods (such as HiCNN) are unable to perform because they require a sparse Hi-C contact map. To test how GrapHiC performs without a Hi-C contact map, we impute GM12878, IMR90, and K562 cell line Hi-C contact maps using cell line-specific ChIP-seq signals and an expected Hi-C contact map. We construct this expected Hi-C contact map using a simple prior that captures the likelihood of observing a Hi-C read, which decays exponentially as the distance between the loci increases. Under the graph structure learning frameworks, we can rely on the expected Hi-C to provide structural topology similar to a sparse Hi-C map coupled with ChIP-seq can provide sufficiently informative representations. We retrain a GrapHiC version that only takes a ChIP-seq and an expected Hi-C contact map to impute GM12878, K562, and IMR90 Hi-C contact maps.

Our quantitative results in Table 2 detail the performance of GrapHiC across all the predicted samples when provided an expected Hi-C contact map. Across the three cell lines, we observe that using an expected Hi-C contact map as a structural prior improves the performance scores on average over HiCReg (a Seq-to-HiC) baseline by a significant 1.4, 2 and 2 times on SSIM, GenomeDISCO, and Chromatin Loops F1 scores. This substantial improvement can be attributed partly to the graph representation we employ in GrapHiC (Graph autoencoder vs. Random Forest in HiCReg) and partly to the value of providing the structural neighborhood information through an expected Hi-C contact map. Conversely, we do observe a reduction in performance on average across all three cell lines by 0.75%, 6.4%, 26% on SSIM, GenomeDISCO, and Chromatin Loops F1 score in comparison to the scenario where we do provide a Hi-C contact map. This reduction in performance can be further bridged by implementing more accurate priors based on known mediators of structural interactions. For instance, genomic loci with high GC content density [4] are more prone to interact, which can improve the prior we provide to GrapHiC. In favor of saving space, we show the visualizations of some selected regions in the Supplementary Fig. S3, which shows that GrapHiC with an expected Hi-C contact map is able to better recover chromatin features in comparison to HiCReg.

**Table 2.** HiCReg, GrapHiC with a Hi-C contact map and GrapHiC without a Hi-C contact map (with a structural prior) scores on SSIM, GenomeDISCO, and Chromatin Loop recall analysis on GM12878, IMR90 and K562. GrapHiC without a Hi-C contact map is able to achieve substantially better than HiCReg a Seq-to-HiC method across all metrics on all three cell lines.

|  | SSIM |  |  | GenomeDISCO |  |  | Chromatin Loops |  |  |
| --- | --- | --- | --- | --- | --- | --- | --- | --- | --- |
|  | HiCReg | GraphHiC-Without HiC | GraphHiC-With HiC | HiCReg | GraphHiC-Without HiC | GraphHiC-With HiC | HiCReg | GraphHiC-Without HiC | GraphHiC-With HiC |
| GM12878 | 0.73768 | 0.90830 | **0.92244** | 0.63565 | 0.84428 | **0.91000** | 0.18041 | 0.36520 | **0.52160** |
| IMR90 | 0.51890 | 0.77460 | **0.78031** | 0.18240 | 0.57363 | **0.61493** | 0.15290 | 0.37280 | **0.47150** |
| K562 | 0.54135 | **0.82800** | 0.82780 | 0.36215 | 0.80279 | **0.84704** | 0.19952 | 0.33050 | **0.45220** |

## 6 Discussion and Conclusion

We present a robust method, GrapHiC, to formulate the Hi-C data as a graph, which is an accurate representation of the chromatin structure. GrapHiC allows us to integrate multiple types of diverse information signals, such as DNA accessibility information, TF binding sites, histone modifications, and spatial arrangement of DNA, into a single representation learning framework. This formulation can be utilized for tasks other than Hi-C read imputation, such as cell-phase identification, cell clustering, chromatin loops, and TAD boundary identification because GrapHiC constructs an information-rich representation.

Our evaluations show that GrapHiC is resilient against the sparsity of the real-world input Hi-C data and can reliably impute Hi-C reads that are biologically informative even in cases when Hi-C experiment data is unavailable. Currently, we are using a simple structural prior; we can improve on that substantially by incorporating the influence of known features such as GC content. GrapHiC can extend the efforts of Avocado [31] to impute missing Hi-C experiments for all the cell lines on the ENCODE portal [21]. We plan to investigate how GrapHiC generalizes to data generated by different Hi-C protocol variations, such as Pore-C [34] and single-cell Hi-C, where the Hi-C reads tend to be very sparse and can potentially develop a bridge between the learning that happens on bulk experiments to the learning we are required to do for single-cell experiments. We plan to tackle these tasks in future investigations as they bring challenges that require data-specific modeling and handling.

## Funding and Acknowledgements

Ghulam Murtaza’s effort on this project was funded by the NIST PREP grant GR5245041. Ritambhara Singh’s contribution to the work is supported by NIH award 1R35HG011939-01. Certain commercial equipment, instruments, or materials are identified to specify adequately experimental conditions or reported results. Such identification does not imply recommendation or endorsement by the National Institute of Standards and Technology, nor does it imply that the equipment, instruments, or materials identified are necessarily the best available for the purpose.

## Footnotes

1 synonymous with low-read-count, low-resolution, or low-quality

2 WL algoritm tries to assign a unique color to each node based on its neighborhood.

3 All of our datasets are aligned with the hg19 reference genome, and we show minor difference (2%) in performance when we provide a grch38 aligned inputs as shown in Supplementary Table. S3.

4 Our GitHub repository contains a link that includes visualizations of all regions of the test chromosomes

5 Integrating different types of datasets or modalities

## References

[1] Tanya Barrett, Stephen E. Wilhite, and et al. 2012. NCBI GEO: Archive for functional genomics data sets—update. Nucleic Acids Research 41, D1 (2012). https://doi.org/10.1093/nar/gks1193

[2] Jeremy Bigness, Xavier Loinaz, and et al. 2022. Integrating longrange regulatory interactions to predict gene expression using graph convolutional networks. Journal of Computational Biology 29, 5 (2022), 409–424. https://doi.org/10.1089/cmb.2021.0316

[3] Kate B Cook, Borislav H Hristov, and et al. 2020. Measuring significant changes in chromatin conformation with accost. Nucleic Acids Research 48, 5 (2020), 2303–2311. https://doi.org/10.1093/nar/gkaa069

[4] Job Dekker. 2007. GC- and AT-rich chromatin domains differ in conformation and histone modification status and are differentially modulated by RPD3P. Genome Biology 8, 6 (2007). https://doi.org/10.1186/gb-2007-8-6-r116

[5] Jesse R. Dixon, Siddarth Selvaraj, Feng Yue, Audrey Kim, Yan Li, Yin Shen, Ming Hu, Jun S. Liu, and Bing Ren. 2012. Topological domains in mammalian genomes identified by analysis of chromatin interactions. Nature 485, 7398 (2012), 376–380. https://doi.org/10.1038/nature11082

[6] Chao Dong, Chen Change Loy, and et al. 2015. Image Super-Resolution Using Deep Convolutional Networks. https://doi.org/10.48550/ARXIV.1501.00092

[7] Vijay Prakash Dwivedi, Chaitanya K. Joshi, and et al. [n. d.]. Benchmarking Graph Neural Networks. https://doi.org/10.48550/ARXIV.2003.00982

[8] Fan Feng, Yuan Yao, Xue Qing David Wang, Xiaotian Zhang, and Jie Liu. 2022. Connecting high-resolution 3D chromatin organization with epigenomics. Nature communications 13, 1 (2022), 1–10.

[9] Alireza Fotuhi Siahpirani, Ferhat Ay, and Sushmita Roy. 2016. A multitask graph-clustering approach for chromosome conformation capture data sets identifies conserved modules of chromosomal interactions. Genome Biology 17, 1 (2016).

[10] Geoff Fudenberg, David R Kelley, and Katherine S Pollard. 2020. Predicting 3D genome folding from DNA sequence with Akita. Nature Methods 17, 11 (2020), 1111–1117.

[11] Max Highsmith and Jianlin Cheng. 2020. Vehicle: A variationally encoded hi-C loss enhancement algorithm. Scientific Reports (2020). https://doi.org/10.1101/2020.12.07.413559

[12] Hao Hong, Shuai Jiang, and et al. 2020. DeepHiC: A generative adversarial network for enhancing hi-C data resolution. PLOS Computational Biology 16, 2 (2020). https://doi.org/10.1371/journal.pcbi.1007287

[13] Dirk A Kleinjan, Anne Seawright, and et al. 2001. Aniridia-associated translocations, DNase hypersensitivity, sequence comparison and transgenic analysis redefine the functional domain of PAX6. Human molecular genetics 10, 19 (2001), 2049–2059.

[14] Simon Kornblith, Ting Chen, Honglak Lee, and Mohammad Norouzi. [n. d.]. Why Do Better Loss Functions Lead to Less Transferable Features? https://doi.org/10.48550/ARXIV.2010.16402

[15] Jack Lanchantin and Yanjun Qi. 2020. Graph convolutional networks for epigenetic state prediction using both sequence and 3D genome data. Bioinformatics 36, Supplement2 (2020), i659–i667. https://doi.org/10.1093/bioinformatics/btaa793

[16] Da-Inn Lee and Sushmita Roy. 2021. Grinch: Simultaneous smoothing and detection of topological units of genome organization from sparse chromatin contact count matrices with matrix factorization. Genome Biology 22, 1 (2021). https://doi.org/10.1186/s13059-021-02378-z

[17] Erez Lieberman-Aiden, Nynke L. van Berkum, Louise Williams, and et al. 2009. Comprehensive Mapping of Long-Range Interactions Reveals Folding Principles of the Human Genome. Science 326, 5950 (2009), 289–293. https://doi.org/10.1126/science.1181369

[18] Qiao Liu, Hairong Lv, and Rui Jiang. 2019. hicGAN infers super resolution Hi-C data with generative adversarial networks. Bioinformatics 35, 14 (2019), i99–i107.

[19] Tong Liu and Zheng Wang. 2019. HiCNN: a very deep convolutional neural network to better enhance the resolution of Hi-C data. Bioinformatics 35, 21 (04 2019), 4222–4228. https://doi.org/10.1093/bioinformatics/btz251

[20] Tong Liu and Zheng Wang. 2019. HiCNN2: Enhancing the Resolution of Hi-C Data Using an Ensemble of Convolutional Neural Networks. Genes 10, 11 (2019). https://doi.org/10.3390/genes10110862

[21] Yunhai Luo, Benjamin C Hitz, and et al. 2019. New Developments on the encyclopedia of DNA elements (encode) Data Portal. Nucleic Acids Research 48, D1 (2019). https://doi.org/10.1093/nar/gkz1062

[22] Cyril Matthey-Doret, Lyam Baudry, Axel Breuer, Remi Montagne, Nadge Guiglielmoni, Vittore Scolari, Etienne Jean, Arnaud Campeas, Philippe Henri Chanut, Edgar Oriol, et al. 2020. Computer vision for pattern detection in chromosome contact maps. Nature communications 11 (2020).

[23] Federico Monti and et al. 2016. Geometric deep learning on graphs and manifolds using mixture model CNNs. https://doi.org/10.48550/ARXIV.1611.08402

[24] Antonio Mora, Geir Kjetil Sandve, and et al. 2015. In the loop: Promoter–enhancer interactions and bioinformatics. Briefings in Bioinformatics (2015). https://doi.org/10.1093/bib/bbv097

[25] Ghulam Murtaza, Atishay Jain, Madeline Hughes, Thulasi Varatharajan, and Ritambhara Singh. 2022. Investigating the performance of deep learning methods for Hi-C resolution improvement. bioRxiv (2022). https://doi.org/10.1101/2022.01.27.477975

[26] Olivier Petit, Nicolas Thome, Clément Rambour, and Luc Soler. 2021. U-Net Transformer: Self and Cross Attention for Medical Image Segmentation. arXiv:2103.06104 [eess.IV]

[27] Elena M. Pugacheva, Naoki Kubo, Dmitri Loukinov, and et al. 2020. CTCF mediates chromatin looping via N-terminal domain-dependent cohesin retention. Proceedings of the National Academy of Sciences 117, 4 (2020), 2020–2031. https://doi.org/10.1073/pnas.1911708117

[28] Ladislav Rampasek, Mikhail Galkin, Vijay Prakash Dwivedi, Anh Tuan Luu, Guy Wolf, and Dominique Beaini. 2022. Recipe for a General, Powerful, Scalable Graph Transformer. Advances in Neural Information Processing Systems 35 (2022).

[29] Rao and et al. 2014. A 3D Map of the Human Genome at Kilobase Resolution Reveals Principles of Chromatin Looping. Cell 159, 7 (2014), 1665–1680. https://doi.org/10.1016/j.cell.2014.11.021

[30] et al Rao S. 2014. A 3D map of the human genome at kilobase resolution reveals principles of chromatin looping. Cell 159, 7 (2014), 1665–1680.

[31] Jacob Schreiber, Timothy Durham, Jeffrey Bilmes, and William Stafford Noble. 2020. Avocado: A multi-scale deep tensor factorization method learns a latent representation of the human epigenome. Genome Biology 21, 1 (2020). https://doi.org/10.1186/s13059-020-01977-6

[32] Yunsheng Shi, Zhengjie Huang, Shikun Feng, Hui Zhong, Wenjin Wang, and Yu Sun. 2020. Masked Label Prediction: Unified Message Passing Model for Semi-Supervised Classification. https://doi.org/10.48550/ARXIV.2009.03509

[33] Jimin Tan, Javier Rodriguez-Hernaez, and et al. 2022. Cell type-specific prediction of 3D chromatin architecture. (2022). https://doi.org/10.1101/2022.03.05.483136

[34] Netha Ulahannan, Matthew Pendleton, Aditya Deshpande, and Et al. 2019. Nanopore sequencing of DNA concatemers reveals higher-order features of chromatin structure. bioRxiv (2019). https://doi.org/10.1186/s13059-019-1658-7

[35] Galip Gürkan Yardımcı and et al. 2019. Measuring the reproducibility and quality of hi-C data - genome biology. https://doi.org/10.1186/s13059-019-1658-7

[36] Shilu Zhang, Chasman, and et al. 2019. In silico prediction of highresolution hi-C interaction matrices. Nature Communications 10, 1 (2019). https://doi.org/10.1038/s41467-019-13423-8

[37] Yan Zhang, Lin An, and et al. 2018. Enhancing Hi-C data resolution with deep convolutional neural network HiCPlus. Nature communications 9, 1 (2018), 1–9.

