## Supplementary Materials for "GrapHiC: An integrative graph based approach for imputing missing Hi-C reads"

| Component | | Layer | Input Shape | Output Shape |
|---|---|---|---|---|
| **Graph Encoder** | | TransformerConv(13, 32, heads=4) | (-1, 256, 13) | (-1, 256, 32) |
| | | Linear(in_features=128, out_features=32, bias=True) | (-1, 256, 32) | (-1, 256, 32) |
| | | GraphNorm(32) | (-1, 256, 32) | (-1, 256, 32) |
| **Graph Decoder** | **InnerProductDecoder** | InnerProduct | (-1, 256, 32) | (-1, 1, 256, 256) |
| | | Conv2d(1, 32, kernel_size=(3, 3), stride=(1, 1), padding=(1, 1)) | (-1, 256, 256) | (-1, 32, 256, 256) |
| | **DownBlock** | GroupNorm(8, 128, eps=1e-05, affine=True) | (-1, 32, 256, 256) | (-1, 32, 256, 256) |
| | | Conv2d(128, 64, kernel_size=(3, 3), stride=(1, 1), padding=(1, 1)) | (-1, 32, 256, 256) | (-1, 32, 256, 256) |
| | | GroupNorm(8, 64, eps=1e-05, affine=True) | (-1, 32, 256, 256) | (-1, 32, 256, 256) |
| | | Conv2d(64, 64, kernel_size=(3, 3), stride=(1, 1), padding=(1, 1)) | (-1, 32, 256, 256) | (-1, 32, 256, 256) |
| | | Conv2d(32, 32, kernel_size=(3, 3), stride=(2, 2), padding=(1, 1)) | (-1, 32, 256, 256) | (-1, 32, 128, 128) |
| | **DownBlock** | GroupNorm(8, 128, eps=1e-05, affine=True) | (-1, 32, 128, 128) | (-1, 32, 128, 128) |
| | | Conv2d(128, 64, kernel_size=(3, 3), stride=(1, 1), padding=(1, 1)) | (-1, 32, 128, 128) | (-1, 32, 128, 128) |
| | | GroupNorm(8, 64, eps=1e-05, affine=True) | (-1, 32, 128, 128) | (-1, 32, 128, 128) |
| | | Conv2d(64, 64, kernel_size=(3, 3), stride=(1, 1), padding=(1, 1)) | (-1, 32, 128, 128) | (-1, 32, 128, 128) |
| | | Conv2d(32, 32, kernel_size=(3, 3), stride=(2, 2), padding=(1, 1)) | (-1, 32, 128, 128) | (-1, 32, 64, 64) |
| | **DownBlock** | GroupNorm(8, 128, eps=1e-05, affine=True) | (-1, 32, 64, 64) | (-1, 32, 64, 64) |
| | | Conv2d(128, 64, kernel_size=(3, 3), stride=(1, 1), padding=(1, 1)) | (-1, 32, 64, 64) | (-1, 32, 64, 64) |
| | | GroupNorm(8, 64, eps=1e-05, affine=True) | (-1, 32, 64, 64) | (-1, 32, 64, 64) |
| | | Conv2d(64, 64, kernel_size=(3, 3), stride=(1, 1), padding=(1, 1)) | (-1, 32, 64, 64) | (-1, 32, 64, 64) |
| | | Conv2d(32, 32, kernel_size=(3, 3), stride=(2, 2), padding=(1, 1)) | (-1, 32, 64, 64) | (-1, 64, 32, 32) |
| | **Middle Block** | SelfAttention() | (-1, 64, 32, 32) | (-1, 64, 32, 32) |
| | **UpBlock** | Concatenate(Downblock + Previous Block) | (-1, 64, 32, 32)*2 | (-1, 128, 32, 32) |
| | | GroupNorm(8, 64, eps=1e-05, affine=True) | (-1, 128, 32, 32) | (-1, 128, 32, 32) |
| | | Conv2d(128, 64, kernel_size=(3, 3), stride=(1, 1), padding=(1, 1)) | (-1, 128, 32, 32) | (-1, 64, 32, 32) |
| | | GroupNorm(8, 64, eps=1e-05, affine=True) | (-1, 64, 32, 32) | (-1, 64, 32, 32) |
| | | Conv2d(64, 32, kernel_size=(3, 3), stride=(1, 1), padding=(1, 1)) | (-1, 64, 32, 32) | (-1, 32, 32, 32) |
| | | SelfAttention() | (-1, 32, 32, 32) | (-1, 32, 32, 32) |
| | | ConvTranspose2d(32, 32, kernel_size=(4, 4), stride=(2, 2), padding=(1, 1)) | (-1, 64, 32, 32) | (-1, 32, 64, 64) |
| | **UpBlock** | Concatenate(Downblock + Previous Block) | (-1, 32, 64, 64)*2 | (-1, 64, 64, 64) |
| | | GroupNorm(8, 64, eps=1e-05, affine=True) | (-1, 64, 64, 64) | (-1, 64, 64, 64) |
| | | Conv2d(64, 32, kernel_size=(3, 3), stride=(1, 1), padding=(1, 1)) | (-1, 64, 64, 64) | (-1, 32, 64, 64) |
| | | GroupNorm(8, 32, eps=1e-05, affine=True) | (-1, 32, 64, 64) | (-1, 32, 64, 64) |
| | | Conv2d(32, 32, kernel_size=(3, 3), stride=(1, 1), padding=(1, 1)) | (-1, 32, 64, 64) | (-1, 32, 64, 64) |
| | | SelfAttention() | (-1, 32, 64, 64) | (-1, 32, 64, 64) |
| | | ConvTranspose2d(32, 32, kernel_size=(4, 4), stride=(2, 2), padding=(1, 1)) | (-1, 32, 64, 64) | (-1, 32, 128, 128) |
| | **UpBlock** | Concatenate(Downblock + Previous Block) | (-1, 32, 128, 128)*2 | (-1, 64, 128, 128) |
| | | GroupNorm(8, 64, eps=1e-05, affine=True) | (-1, 64, 128, 128) | (-1, 64, 128, 128) |
| | | Conv2d(64, 32, kernel_size=(3, 3), stride=(1, 1), padding=(1, 1)) | (-1, 64, 128, 128) | (-1, 32, 128, 128) |
| | | GroupNorm(8, 32, eps=1e-05, affine=True) | (-1, 32, 128, 128) | (-1, 32, 128, 128) |
| | | Conv2d(32, 32, kernel_size=(3, 3), stride=(1, 1), padding=(1, 1)) | (-1, 32, 128, 128) | (-1, 32, 128, 128) |
| | | SelfAttention() | (-1, 32, 128, 128) | (-1, 32, 128, 128) |
| | | ConvTranspose2d(32, 32, kernel_size=(4, 4), stride=(2, 2), padding=(1, 1)) | (-1, 32, 128, 128) | (-1, 32, 256, 256) |
| | **Final Projection** | GroupNorm(8, 32, eps=1e-05, affine=True) | (-1, 32, 256, 256) | (-1, 32, 256, 256) |
| | | Conv2d(32, 1, kernel_size=(3, 3), stride=(1, 1), padding=(1, 1)) | (-1, 32, 256, 256) | (-1, 1, 256, 256) |
| | | Sigmoid() | (-1, 1, 256, 256) | (-1, 1, 256, 256) |

**Table S1.** This table provides the detailed breakdown of all the layers in our GrapHiC model.

| | Reads | Sparsity | Source |
|---|---|---|---|
| **GRCh38-GM12878-HRC** | 6,524,520,477 | 1 | ENCFF555ISR |
| **GRCh38-GM12878-LRC** | 283,697,048 | 23 | ENCFF216ZNY |
| **GRCh38-K562-HRC** | 2,188,905,398 | 1 | ENCFF080DPJ |
| **GRCh38-K562-LRC** | 608,231,511 | 4 | 4DNESI7DEJTM |

**Table S2.** We add four Hi-C datasets, two from GM12878 and two from K562 that are aligned to the GRCh38 assembly to evaluate how GrapHiC generalizes to different assemblies.

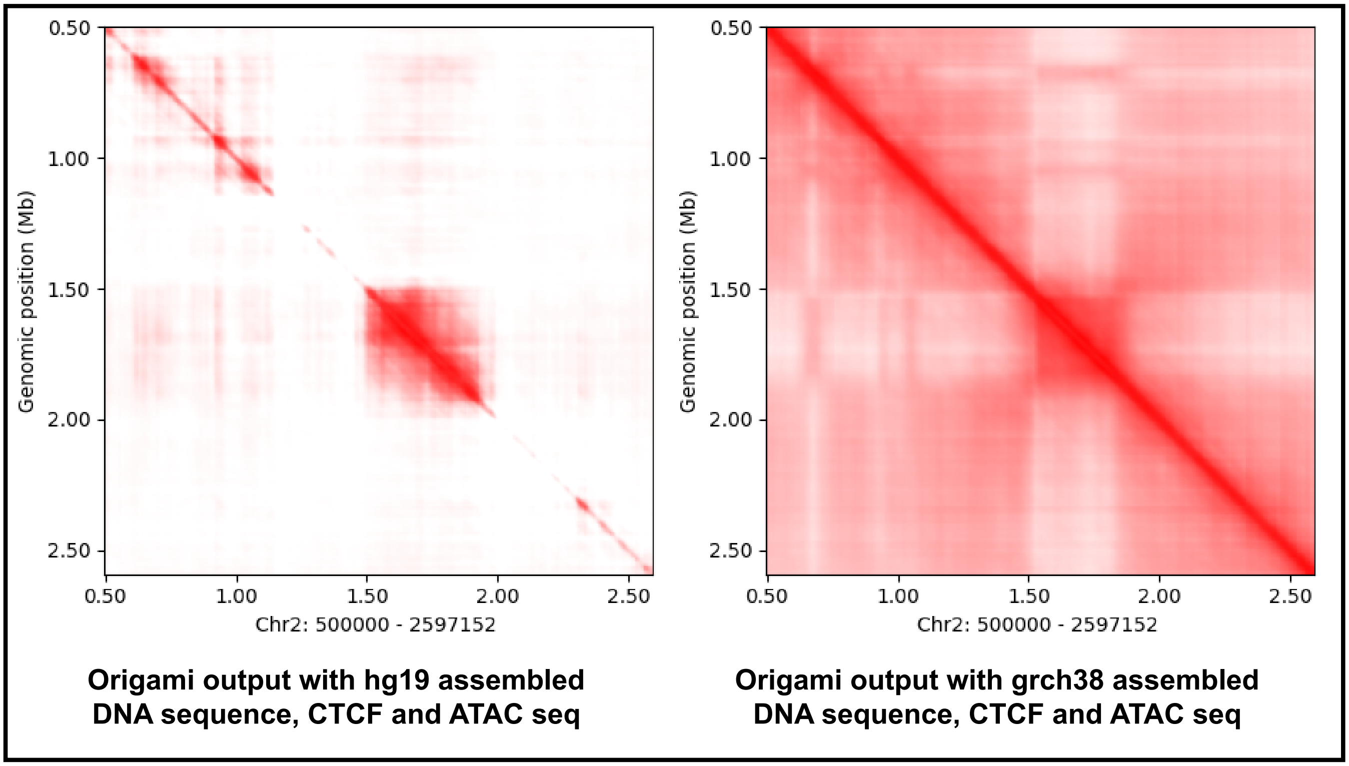


**Figure S1.** We compare the output of Origami, with inputs aligned on hg19 genome assembly against the grch38 aligned assembly. Origami does not generalize to hg19 assembled inputs and struggles to produce meaningful contact maps.

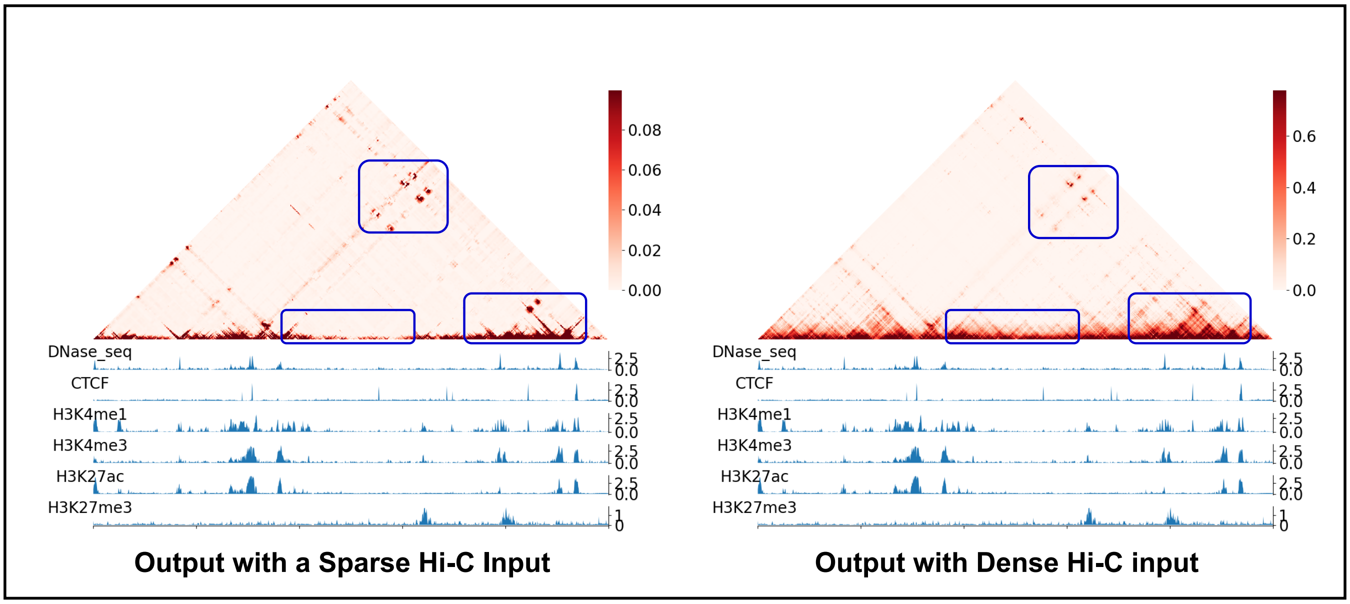


**Figure S2.** We compare the output of Caesar when provided with a sparse real-world H1 cell line Hi-C contact map as input. Caesar struggles to recover distal and nearby features to the diagonal. We provided the same ChIP-seq inputs in both cases. Moreover, Caesar produces MicroC contact maps showing substantially different read contact distributions compared to Hi-C, so we exclude Caesar from our baselines.

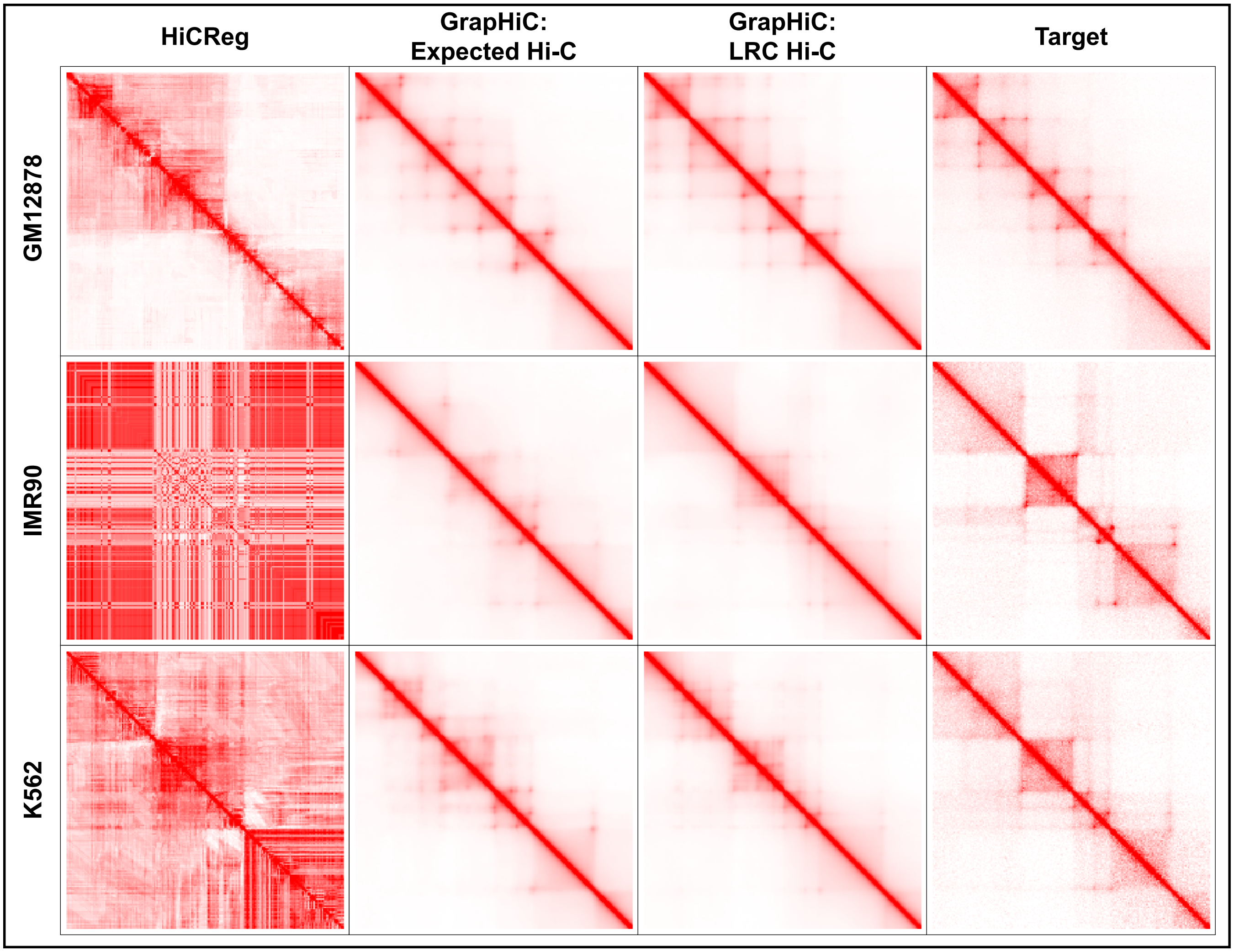


**Figure S3.** We qualitatively compare the output of GrapHiC when provided with a expected Hi-C contact map, a low-read-count (LRC) contact map against the target and HiCReg. We show that GrapHiC is able to impute high-fidelity Hi-C contact maps in both cases that are more simlar to the target in comparison to HiCReg.

| | | MSE | SSIM | PCC | HiCRep | GenomeDISCO | HiCSpector | QuASAR-Rep | TAD Boundaries | Chromatin Loops | DNA Hairpins |
|---|---|---|---|---|---|---|---|---|---|---|---|
| GM12878 | hg19 | **0.0009** | **0.9224** | **0.9797** | **0.8008** | **0.9100** | **0.4614** | **0.8609** | **0.6648** | **0.5216** | **0.5272** |
| | grch38 | 0.0016 | 0.9053 | 0.9544 | 0.7933 | 0.8822 | 0.4367 | 0.8699 | 0.6514 | 0.5119 | 0.4946 |
| K562 | hg19 | 0.0047 | 0.8278 | 0.8731 | 0.7088 | **0.8470** | **0.3252** | 0.7342 | 0.5388 | **0.4522** | 0.3703 |
| | grch38 | **0.0024** | **0.8747** | **0.9262** | **0.7127** | 0.7753 | 0.3152 | **0.7848** | **0.5698** | 0.4386 | **0.4297** |

**Table S3.** We compare the performance of GrapHiC trained on hg19 aligned datasets on Hg19 and GRCh38 aligned datasets for GM12878 and K562 cell lines. We observe a minor decrease in the GM12878 cell line scores; we believe this change arises because the GRCh38 GM12878 HRC dataset has a substantially higher number of reads in comparison to Hg19 GM12878 HRC (1.8 billion vs. 6.4 billion reads). The Hi-C contact maps generated by GrapHiC match the feature distribution of the 1.8 billion reads contact map and have a smaller set of features compared to the 6.4 billion reads contact map. This difference manifests as a degradation in scores. Conversely, we observe an improvement in scores on the K562 dataset because now the GRCh38 K562 Hi-C dataset has a sequencing depth more similar to the number of reads in the Hg19 Hi-C contact map (1.9 billion vs. 2.2 billion), and this similarity of feature density in both contact maps manifests as improvement in scores. There are distributional differences in both Hg19 assembled Hi-C contact maps, and GRCh38 assembled contact maps that we plan to investigate in more detail as part of our future work. We have also released the GrapHiC weights trained for GRCh38 model available.

| | MSE | SSIM | PCC | HiCRep | GenomeDISCO | HiCSpector | QuASAR-Rep | TAD Boundaries | Chromatin Loops | DNA Hairpins |
|---|---|---|---|---|---|---|---|---|---|---|
| **graphic-basic** | 0.00181278963 | 0.838731225 | 0.9556718263 | 0.3561238127 | 0.6889723173 | 0.2561238791 | 0.6187238192 | 0.456128371 | 0.1251243891 | 0.101512812 |
| **graphic-pos** | 0.001040138886 | 0.9118280711 | 0.9744101287 | 0.8011612903 | 0.8011612903 | 0.4349677419 | 0.8416831613 | 0.599158595 | 0.4393064431 | 0.4483152259 |
| **graphic-ctcf** | 0.000885409594 | 0.923217525 | 0.9793579727 | 0.831483871 | 0.8572446452 | **0.4888709677** | 0.8572446452 | 0.6730900735 | 0.5416144993 | 0.5285637621 |
| **graphic** | **0.0008290408296** | **0.9239148305** | **0.9805694712** | 0.8364516129 | **0.9089642857** | 0.4707096774 | 0.8614972581 | 0.6757557069 | **0.5484932834** | **0.5475020008** |
| **graphic-large** | 0.0008437751676 | 0.9226424879 | 0.9802531959 | **0.8576774194** | 0.8943636364 | 0.4867419355 | **0.8756502903** | **0.7003308268** | 0.5374315339 | 0.5385440433 |

**Table S4.** This table shows detailed ablations results on the GM12878-LRC-1 Hi-C dataset.

| | MSE | SSIM | PCC | HiCRep | GenomeDISCO | HiCSpector | QuASAR-Rep | TAD Boundaries | Chromatin Loops | DNA Hairpins |
|---|---|---|---|---|---|---|---|---|---|---|
| **graphic-basic** | 0.00171278963 | 0.8461231251 | 0.955657121 | 0.3912381273 | 0.7081230912 | 0.263891273 | 0.6571283012 | 0.4981237819 | 0.2025871212 | 0.179289931 |
| **graphic-pos** | 0.0009828922339 | 0.9128129021 | 0.9760758355 | 0.783516129 | 0.8600357143 | 0.4607419355 | 0.8362809677 | 0.6173654048 | 0.4367204234 | 0.4443665519 |
| **graphic-ctcf** | 0.0008680545725 | 0.9223760309 | 0.9798637377 | 0.7907096774 | 0.8942413793 | **0.4751612903** | 0.8521077742 | 0.6845039088 | 0.5351110329 | 0.5324476257 |
| **graphic** | **0.0008289036923** | **0.9228198434** | **0.9805673792** | 0.8179032258 | **0.9030384615** | 0.457 | 0.8575350323 | 0.6821782829 | **0.5365461676** | **0.5487044283** |
| **graphic-large** | 0.000847046962 | 0.922221218 | 0.9803199076 | **0.8322580645** | 0.8888636364 | 0.4499032258 | **0.8671765161** | **0.6968594479** | 0.5194946749 | 0.5372154113 |

**Table S5.** This table shows detailed ablations results on the GM12878-LRC-2 Hi-C dataset.

| | MSE | SSIM | PCC | HiCRep | GenomeDISCO | HiCSpector | QuASAR-Rep | TAD Boundaries | Chromatin Loops | DNA Hairpins |
|---|---|---|---|---|---|---|---|---|---|---|
| **graphic-basic** | 0.0017591283 | 0.8451236123 | 0.9563819028 | 0.4015812312 | 0.7123019823 | 0.268317541 | 0.6667238192 | 0.5123871251 | 0.2312873196 | 0.191512812 |
| **graphic-pos** | 0.0009522660403 | 0.9186408093 | 0.9774438178 | 0.8046774194 | 0.8921034483 | 0.4806129032 | 0.8345150968 | 0.5999143664 | 0.4356771671 | 0.39638861 |
| **graphic-ctcf** | 0.0009 | 0.9222859577 | 0.9793420255 | 0.8116451613 | 0.9049285714 | **0.4852903226** | 0.8512654516 | 0.6683 | 0.5164511455 | 0.5167336452 |
| **graphic** | 0.0008672421682 | 0.9224354332 | 0.9797065909 | 0.8007741935 | **0.91** | 0.4613548387 | 0.8608621613 | 0.6647960639 | **0.5216039742** | **0.52716** |
| **graphic-large** | **0.0008528126054** | **0.9224282913** | **0.9801922214** | **0.8458709677** | 0.8855833333 | 0.4707741935 | **0.8617122903** | **0.6885320461** | 0.5096508442 | 0.525595418 |

**Table S6.** This table shows detailed ablations results on the GM12878-LRC-3 Hi-C dataset.

| | MSE | SSIM | PCC | HiCRep | GenomeDISCO | HiCSpector | QuASAR-Rep | TAD Boundaries | Chromatin Loops | DNA Hairpins |
|---|---|---|---|---|---|---|---|---|---|---|
| **graphic-basic** | 0.00171278591 | 0.8356182415 | 0.9551293856 | 0.3981293789 | 0.7032179831 | 0.2695812031 | 0.6538921731 | 0.5021987451 | 0.2212873126 | 0.171512812 |
| **graphic-pos** | 0.0009809146868 | 0.9193714093 | 0.9764888496 | 0.7997096774 | 0.8847586207 | 0.4398064516 | 0.838564129 | 0.5726642243 | 0.3868603552 | 0.4073110319 |
| **graphic-ctcf** | 0.0009762486443 | 0.9200183289 | 0.9776983314 | 0.8104193548 | 0.8972068966 | **0.4779354839** | 0.8548853548 | 0.6748089642 | 0.5089955344 | 0.5018595278 |
| **graphic** | 0.0009478544234 | **0.920667914** | 0.9779336097 | 0.8148064516 | **0.9058461538** | 0.4660322581 | 0.8636007333 | 0.6639213627 | **0.5150424406** | **0.5337003302** |
| **graphic-large** | **0.0008896560175** | 0.9204220266 | **0.9792669186** | **0.8453225806** | 0.88308 | 0.4606451613 | **0.8677272903** | **0.6812338361** | 0.4997126192 | 0.5250535956 |

**Table S7.** This table shows detailed ablations results on the GM12878-LRC-4 Hi-C dataset.

| | MSE | SSIM | PCC | HiCRep | GenomeDISCO | HiCSpector | QuASAR-Rep | TAD Boundaries | Chromatin Loops | DNA Hairpins |
|---|---|---|---|---|---|---|---|---|---|---|
| **graphic-basic** | 0.00201278963 | 0.828128741 | 0.9467128312 | 0.3017892319 | 0.625198231 | 0.2561238791 | 0.5517238112 | 0.430897451 | 0.1181023712 | 0.101512812 |
| **graphic-pos** | 0.001829135232 | 0.8873864556 | 0.9519068669 | 0.4235806452 | 0.7322068966 | 0.276516129 | 0.7074913548 | 0.3248079352 | 0.155861061 | 0.1206382959 |
| **graphic-ctcf** | 0.001466627116 | 0.8959276496 | 0.9661399373 | 0.5114516129 | 0.75724 | 0.2982258065 | 0.7384491667 | 0.4848169158 | 0.3012013986 | 0.3201851551 |
| **graphic** | 0.001204534899 | 0.9005296682 | 0.9698582723 | 0.5322903226 | 0.8051363636 | 0.3179677419 | 0.7461324138 | 0.5386603403 | 0.3410283165 | 0.3874506211 |
| **graphic-large** | **0.001106260577** | **0.902333818** | **0.9728707875** | **0.5829677419** | **0.8374375** | **0.3242903226** | **0.7815497333** | **0.5535101466** | **0.3868443661** | **0.4281288537** |

**Table S8.** This table shows detailed ablations results on the GM12878-LRC-5 Hi-C dataset.

| | | MSE | SSIM | PCC | HiCRep | GenomeDISCO | HiCSpector | QuASAR-Rep | TAD Boundaries | Chromatin Loops | DNA Hairpins |
|---|---|---|---|---|---|---|---|---|---|---|---|
| **GM12878-LRC-1** | **HiCReg** | 0.00525 | 0.73768 | 0.86519 | 0.37990 | 0.63565 | 0.24605 | 0.68039 | 0.41440 | 0.18041 | 0.22014 |
| | **HiCNN** | 0.00150 | 0.90890 | 0.97770 | **0.90182** | 0.80220 | 0.46291 | **0.87124** | **0.71041** | 0.53060 | 0.52280 |
| | **GrapHiC** | **0.00083** | **0.92391** | **0.98057** | 0.83645 | **0.90896** | **0.47071** | 0.86150 | 0.67576 | **0.54849** | **0.54750** |
| **GM12878-LRC-2** | **HiCReg** | 0.00525 | 0.73768 | 0.86519 | 0.37990 | 0.63565 | 0.24605 | 0.68039 | 0.41440 | 0.18041 | 0.22014 |
| | **HiCNN** | 0.00135 | 0.92020 | 0.97455 | 0.80125 | **0.91780** | 0.45215 | 0.00000 | **0.71103** | 0.51860 | 0.54050 |
| | **GrapHiC** | **0.00083** | **0.92282** | **0.98057** | **0.81790** | 0.90304 | **0.45700** | **0.85754** | 0.68218 | **0.53655** | **0.54870** |
| **GM12878-LRC-3** | **HiCReg** | 0.00525 | 0.73768 | 0.86519 | 0.37990 | 0.63565 | 0.24605 | 0.68039 | 0.41440 | 0.18041 | 0.22014 |
| | **HiCNN** | 0.00414 | 0.83110 | 0.92730 | 0.79512 | 0.82610 | 0.45613 | 0.00000 | **0.72110** | 0.52150 | 0.51940 |
| | **GrapHiC** | **0.00087** | **0.92244** | **0.97971** | **0.80077** | **0.91000** | **0.46135** | **0.86086** | 0.66480 | **0.52160** | **0.52716** |
| **GM12878-LRC-4** | **HiCReg** | 0.00525 | 0.73768 | 0.86519 | 0.37990 | 0.63565 | 0.24605 | 0.68039 | 0.41440 | 0.18041 | 0.22014 |
| | **HiCNN** | 0.00456 | 0.81420 | 0.91770 | 0.79124 | 0.79340 | 0.46215 | 0.00000 | **0.70990** | 0.50560 | 0.45070 |
| | **GrapHiC** | **0.00095** | **0.92067** | **0.97793** | **0.81481** | **0.90585** | **0.46603** | **0.86360** | 0.66392 | **0.51504** | **0.53370** |
| **GM12878-LRC-5** | **HiCReg** | 0.00525 | 0.73768 | 0.86519 | 0.37990 | 0.63565 | 0.24605 | 0.68039 | 0.41440 | 0.18041 | 0.22014 |
| | **HiCNN** | 0.00809 | 0.68010 | 0.83977 | 0.47193 | 0.47380 | 0.28129 | 0.00000 | 0.43825 | 0.22160 | 0.15606 |
| | **GrapHiC** | **0.00120** | **0.90053** | **0.96986** | **0.53229** | **0.80514** | **0.31797** | **0.74613** | **0.53866** | **0.34103** | **0.38745** |

**Table S9.** We show the performance of GrapHiC when provided with five different sparse GM12878 datasets. We compare the performance of GrapHiC against HiCReg, HiCNN and bold score of the best performing method.

| | | MSE | SSIM | PCC | HiCRep | GenomeDISCO | HiCSpector | QuASAR-Rep | TAD Boundaries | Chromatin Loops | DNA Hairpins |
|---|---|---|---|---|---|---|---|---|---|---|---|
| IMR90 | **HiCReg** | 0.05150 | 0.51890 | 0.26980 | 0.13090 | 0.18240 | 0.10670 | 0.17740 | 0.35420 | 0.15290 | 0.08620 |
| | **HiCNN** | **0.00510** | 0.65370 | 0.56760 | **0.80790** | 0.51510 | 0.24840 | 0.00000 | **0.56030** | 0.29290 | 0.26730 |
| | **GrapHiC** | 0.00750 | **0.78031** | **0.80270** | 0.69055 | **0.61493** | **0.28152** | **0.77327** | 0.55940 | **0.47150** | **0.47480** |
| K562 | **HiCReg** | 0.05656 | 0.54135 | 0.48524 | 0.27575 | 0.36215 | 0.15900 | 0.51221 | 0.42812 | 0.19952 | 0.27708 |
| | **HiCNN** | 0.01100 | 0.64760 | 0.61870 | **0.87120** | 0.77460 | 0.29510 | 0.00000 | **0.58190** | 0.35720 | 0.26500 |
| | **GrapHiC** | **0.00470** | **0.82780** | **0.87310** | 0.70884 | **0.84704** | **0.32516** | **0.73423** | 0.53880 | **0.45220** | **0.37030** |

**Table S10.** We show the performance of GrapHiC when provided with two different cell line datasets. We compare the performance of GrapHiC against HiCReg, HiCNN and bold score of the best performing method.

| | MSE | SSIM | PCC | HiCRep | GenomeDISCO | HiCSpector | QuASAR-Rep | TAD Boundaries | Chromatin Loops | DNA Hairpins |
|---|---|---|---|---|---|---|---|---|---|---|
| **GM12878** | 0.00110 | 0.90830 | 0.97220 | 0.59400 | 0.84428 | 0.33185 | 0.77946 | 0.61640 | 0.36520 | 0.44350 |
| **IMR90** | 0.00490 | 0.77460 | 0.85050 | 0.35865 | 0.57363 | 0.24410 | 0.63696 | 0.54500 | 0.37280 | 0.40060 |
| **K562** | 0.00440 | 0.82800 | 0.87410 | 0.47852 | 0.80279 | 0.28448 | 0.63149 | 0.51250 | 0.33050 | 0.31200 |

**Table S11.** We show the performance of GrapHiC when provided with an expected Hi-C contact map across three different cell lines.
